# Lumbosacral spinal cord functional connectivity at rest: From feasibility to reliability

**DOI:** 10.1101/2023.12.12.571073

**Authors:** Ilaria Ricchi, Nawal Kinany, Dimitri Van De Ville

**Affiliations:** Neuro-X Institute, Ecole Polytechnique Fédérale de Lausanne (EPFL), Geneva, 1202, Switzerland; Department of Radiology and Medical Informatics, University of Geneva, Geneva, 1211, Switzerland

## Abstract

In the past decade, exploration of spontaneous blood-oxygen-level-dependent (BOLD) signal fluctuations has expanded beyond the brain to include the spinal cord. While most studies have predominantly focused on the cervical region, the lumbosacral segments play a crucial role in motor control and sensory processing of the lower limbs. Addressing this gap, the aims of the current study were two-fold: first, confirming the presence and nature of organized spontaneous BOLD signals in the human lumbosacral spinal cord; second, systematically assessing the impact of various denoising strategies on signal quality and functional connectivity (FC) patterns. Given the susceptibility of spinal cord fMRI to noise, this step is pivotal to ensure the robustness of intrinsic FC. Our findings uncovered bilateral FC between the ventral horns. Importantly, these patterns were consistently observed across denoising methods and demonstrating fair to excellent reliability. Conversely, no other significant connectivity patterns were identified across the remaining horns. Importantly, the evaluation of diverse denoising strategies highlighted the efficacy of PNM-based pipelines in cleaning the signal while preserving the strength and reliability of connectivity estimates. Together, our results provide evidence of robust FC patterns in the lumbosacral spinal cord, thereby paving the way for future studies probing caudal spinal activity.

## 1. Introduction

Functional magnetic resonance imaging (fMRI) is a non-invasive imaging technique that has revolutionized our ability to investigate the central nervous system (CNS). By exploiting the blood-oxygenation-level-dependent (BOLD) signal, a hemodynamic proxy of neural activity (Logothetis, 2003), fMRI has become a method of choice for exploring brain function. Notably, the acquisition of fMRI data during resting state – marked by the absence of explicit tasks or stimuli – has garnered significant attention. This interest stems from the compelling observation that spontaneous BOLD fluctuations can be parsed into so-called resting-state networks (Biswal et al., 2010; Damoiseaux et al., 2006; Fox and Raichle, 2007), which reflect the brain’s intrinsic functional organization. These networks have been shown to be behaviorally relevant, making them valuable tools to study healthy and impaired brain function (van den Heuvel and Hulshoff Pol, 2010).

More recently, the scope of resting-state fMRI has expanded beyond the confines of the cortex, for instance to probe the intrinsic organization of the spinal cord; see (Harrison et al., 2021) for review. Notably, studies focusing on the cervical spinal cord have uncovered organized spontaneous signals, utilizing both data-driven methods, such as ICA (Kong et al., 2014; Landelle et al., 2021) and iCAP (Kinany et al., 2020)), and hypothesis-driven approaches (Barry et al., 2014; Eippert et al., 2017b; Harita and Stroman, 2017; Kaptan et al., 2023; Liu et al., 2016; Weber et al., 2018). These studies have effectively revealed spinal resting-state networks, prominently featuring functional connectivity between bilateral ventral (*i.e.*, motor) and dorsal (*i.e.*, sensory) horns. Building on these auspicious results, further studies have then explored the reliability of these functional connectivity patterns, a critical consideration to ensure their broader applicability in fundamental and clinical applications. For instance, Barry and colleagues demonstrated their stability within the same scanning session (Barry et al., 2016). Additional investigations have evaluated the impact of different acquisition and processing choices (Barry et al., 2018; Eippert et al., 2017b; Kinany, 2022), as well as the influence of distinct noise sources (Kaptan et al., 2023). These collective efforts have underscored the robustness of functional connectivity patterns in the cervical spinal cord.

In spite of these promising findings, areas situated caudal to the cervical spinal cord have so far largely eluded exploration. Remarkably, the lumbosacral spinal cord, crucial for motor control and sensory processing of the lower limbs, has been largely overlooked. This oversight can be attributed, in part, to the challenges associated with fMRI acquisition, processing, and analysis in this region, stemming from the smaller size of the cord (Frostell et al., 2016), heightened anatomical variability (Tins and Balain, 2016; Van Schoor et al., 2015), and the lack of dedicated tools have compounded this difficulty. To date, only one study (Combes et al., 2023) has deployed resting-state fMRI to uncover the intrinsic organization of the lumbosacral spinal cord, shedding light on sensorimotor networks reminiscent of those observed in the cervical spinal cord. Building on these auspicious observations, a critical aspect yet to be explored pertains to the reliability of these lumbosacral functional connectivity patterns. Addressing this is pivotal for the advancement of resting-state metrics as potent tools to identify functional biomarkers, especially in the context of neurological conditions such as movement disorders and spinal cord injuries (Conrad et al., 2018; Kreiter et al., 2022; Rowald et al., 2022).

To address this knowledge gap, this study sets out to systematically investigate spontaneous BOLD fluctuations in the human lumbosacral spinal cord. Recognizing the susceptibility of spinal cord fMRI to a variety of noise sources, our aims are twofold: firstly, to confirm the presence and nature of organized fluctuations in the lumbosacral spinal cord, employing a sequence distinct from the one used by (Combes et al., 2023); and secondly, to assess the reliability of functional connectivity measures to variations in the denoising procedure. By identifying the optimal approach for assessing functional connectivity in the lumbosacral spinal cord, we intend to contribute to the development of robust methods for studying the functional architecture of this region. Ultimately, our work will enhance our ability to investigate the CNS on a larger scale, opening up new avenues for research and clinical applications.

## 2. Methods

### 2.1. Participants

22 healthy volunteers were enrolled in this study (11 male, 11 female, 27.35 ± 2.34 years old). Participants reported no history of neurological or motor disorders. All participants gave their written informed consent to participate, and the study was approved by the Commission Cantonale d’Éthique de la Recherche Genève (CCER, study 2019-00203).

### 2.2. Data acquisition

All experiments were performed on a Siemens Prisma scanner (3 Tesla) (Erlangen, Germany), equipped with a 32-channel spine coil of which 16 were used for the acquisitions. Participants were placed in the scanner in supine position. Functional images were acquired using a T2*-weighted echo-planar imaging (EPI) sequence with ZOOMit selective field-of-view imaging, based on our previous cervical protocols (Kinany et al., 2022, 2019, 2020) but adapted for the lumbosacral spinal cord (Repetition Time (TR) = 2.5 s, Echo Time (TE) = 34 ms, FOV = 44 × 144, flip angle = 80°, GRAPPA acceleration factor: 2, in-plane resolution = 1.1 × 1.1 mm^2^, slice thickness = 3 mm). Compared to cervical recordings, a wider field-of-view (*i.e.*, changing in-plane resolution from 1 to 1.1 mm) was employed to account for the additional tissue volume present at the levels of the hips, and to avoid aliasing artifacts. The lumbosacral enlargement (approximately from at vertebral levels T11 to L2) was covered using 27 axial slices, positioned perpendicularly to the spinal cord to limit signal dropouts due to field inhomogeneities (Finsterbusch et al., 2012). Manual shimming adjustments focused on the spinal cord were conducted prior to the functional acquisitions to optimize the magnetic field homogeneity. For each participant, 360 volumes (*i.e.*, 15 minutes) were acquired, during rest (*i.e.*, no explicit task) with eyes open (an empty screen was shown). Additionally, a high resolution T2-weighted images (64 sagittal slices; resolution: 0.4 × 0.4 × 0.8mm^3^; field-of-view: 250 × 250mm^2^; TE: 133ms; flip angle: 140^୦^; TR: 1500ms; GRAPPA acceleration factor: 2; partial Fourier factor: 6/8; acquisition time: 6:03 min) was acquired, although it was not used for this study.

During the fMRI data acquisition, we recorded peripheral physiological signals to perform physiological noise modeling: cardiac data were acquired using a photoplethysmograph and respiratory signals were obtained with a belt (Biopac MP150 system, California, USA). Simultaneous recordings of scanner triggers ensured synchronization of the recordings.

### 2.3. Data preprocessing

Preprocessing steps were performed using Python (version 3.9,7), with nilearn library (version 0.9.1) falling under the umbrella of scikit-learn (version 0.24.2), FMRIB Software Library (FSL; version 5.0) and Spinal Cord Toolbox (SCT; version 5.3.0; (De Leener et al., 2017)).

#### 2.3.1. Preprocessing of fMRI data

##### 2.3.1.1. Motion correction

Given the small size of the spinal cord, in particular at the lumbosacral levels (Frostell et al., 2016), motion correction is a crucial step. The volumes of each functional run were averaged and the centerline of the spinal cord was automatically extracted from the resulting image. A cylindrical mask along this centerline was drawn (30 mm) and further used to exclude regions outside the spinal cord, thus limiting the impact of regions that might move independently of the cord. To account for the articulated structure of the spine, slice-wise realignment was then performed using the SCT function *‘sct_fmri_moco’* (De Leener et al., 2017), with no z-regularization and a B-spline interpolation. The amount of motion was assessed by computing the average absolute value of the framewise displacement (FD). The motion correction parameters were afterwards used as regressors for denoising strategy (see section 2.3.2.).

##### 2.3.1.2. Segmentation

For the functional runs, we use the FSLeyes software to manually create binary masks of the spinal cord only and the spinal cord with the surrounding subarachnoid cavity, using mean motion-corrected images. The subtraction of the former from the latter generated the mask of the CSF only, which was manually inspected for each participant.

#### 2.3.2 Denoising and temporal filtering

We systematically investigated the impact of distinct denoising procedures in the lumbosacral spinal cord by applying denoising pipelines incorporating different confounds. A temporal band-pass filter (cut-off frequencies: 0.01 Hz and 0.13 Hz) was applied. Each denoising procedure relies on the usage of the *‘clean_img*’ function from nilearn library, which allows us to remove the noise confounds orthogonally to the temporal filter. Specifically, confounds and temporal filter were projected onto the same orthogonal space, following the methodology outlined in (Lindquist et al., 2019), instead of being applied sequentially.

Physiological data were used to build nuisance regressors, using a model-based approach derived from the RETROspective Image CORrection (RETROICOR) procedure (Glover et al., 2000). This model assumes the physiological signals to be quasi-periodic, which leads to uniquely assigning the cardiac and respiratory phases to each image using a Fourier expansion. To this aim, we resorted to FSL’s physiological noise modeling (PNM) tool to generate regressors from cardiac, respiratory and CSF signals. Cardiac peaks were automatically detected using the *‘scipy.signal.find_peaks’* function (Virtanen et al., 2020), with manual inspection to ensure reliability.

We followed recommendations for PNM in the spinal cord (Kong et al., 2012). For both cardiac and respiratory regressors, we employed an order of 4, which means that the base frequency was used along with the first 3 harmonics. Cardiac and respiratory signals were combined with an interaction order of 2. This resulted in a total of 32 regressors. A CSF regressor was also calculated as the mean signal from the top 10% of CSF voxels with the most signal variability. Of note, the PNM tool generates slice-wise regressors, while the *‘clean_img*’ function expects volume-wise regressors. To align with this requirement, we averaged slice-wise regressors to obtain volume-wise regressors for input into the denoising step.

In addition to model-based denoising approaches, we leveraged the CSF signal to account for non-neural fluctuations using a data-driven method. Specifically, we used a component-based noise correction technique known as CompCor (Behzadi et al., 2007), which estimates K regressors (K set to 5 in our case) corresponding to the most significant principal components derived from CSF noise. We implemented this method using the ‘*nipype.algorithms.confounds’* module. The combination of all the above mentioned procedures resulted in 12 denoising pipelines illustrated in Table 1.

**Table 1.** Denoising pipelines. This table depicts the nuisance regressors taken into account (columns) for each denoising pipeline (rows), along with the count of regressors of interest per option (last row) and the overall total considering the several combinations (last column).

|  | Baseline | CSF | Motion parameters | CompCor | Cardiac | Respiratory | Card-resp interaction | # regressors (total) |
| --- | --- | --- | --- | --- | --- | --- | --- | --- |
| Baseline |  |  |  |  |  |  |  | 0 |
| CSF |  | ✓ |  |  |  |  |  | 1 |
| Moco |  |  | ✓ |  |  |  |  | 2 |
| CompCor |  |  |  | ✓ |  |  |  | 5 |
| Moco + CompCor |  |  | ✓ | ✓ |  |  |  | 7 |
| Cardiac |  |  |  |  | ✓ |  |  | 8 |
| Respiratory |  |  |  |  |  | ✓ |  | 8 |
| PNM |  |  |  |  | ✓ | ✓ | ✓ | 32 |
| PNM + CSF |  | ✓ |  |  | ✓ | ✓ | ✓ | 33 |
| PNM + Moco |  |  | ✓ |  | ✓ | ✓ | ✓ | 34 |
| PNM + Moco + CSF |  | ✓ | ✓ |  | ✓ | ✓ | ✓ | 35 |
| All |  | ✓ | ✓ | ✓ | ✓ | ✓ | ✓ | 40 |
| # regressors (per option) | 0 | 1 | 2 | 5 | 8 | 8 | 16 |  |

#### 2.3.3 Temporal SNR (tSNR) and explained variance

To evaluate the impact of the different denoising procedures on the signals, we calculated the temporal signal-to-noise ratio (tSNR) and the explained variance of the time series within the spinal cord mask. The voxelwise tSNR values were obtained with the SCT’s function *‘sct_fmri_compute_tsnr’* (De Leener et al., 2017) which computes each voxel’s temporal mean and divides it by its standard deviation. TSNR values were also averaged in the four gray matter ROIs. The explained variance (R^2^) was computed as the fractional reduction of signal variance (Birn et al., 2014):

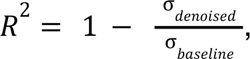

where σ_denoised_ is the variance of the denoised signals for a specific denoising procedure and σ_baseline_ indicates the baseline variance of the time series before denoising (i.e., after motion-correction). The R^2^ values were then adjusted to take into account the number of regressors used in the denoising procedure:

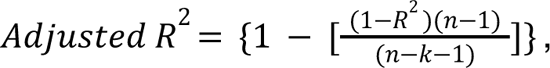

where *n* represents the number of time points and *k* the number of regressors in the nuisance model.

As the Kolmogorov-Smirnov test rejected the normality assumption for the tSNR and R^2^ distributions, we resorted to the Welch’s t-test, which is robust for small sample size and non-normality (Ahad and Syed-Yahaya, 2014). In particular, we compared the explained variance distributions in order of increasing complexity: comparing each denoising method against the previous, less complex, strategy (namely, Moco with CSF, CompCor with Moco, etc.). As for the tSNR, we also relied on the Welch’s t-test, this time to compare each denoising procedure distribution with respect to the baseline.

### 2.4. Data analyses

Functional connectivity (FC) analyses were performed in the native space, using a seed-based approach. Specifically, seeds were manually placed on the slices of the mean functional images, in four specific locations (ventral and dorsal horns on both sides, Fig. 1a). On average, participants had approximately 13 slices in which the gray matter was sufficiently visible to enable confident placement of the seeds.

**Figure 1.**
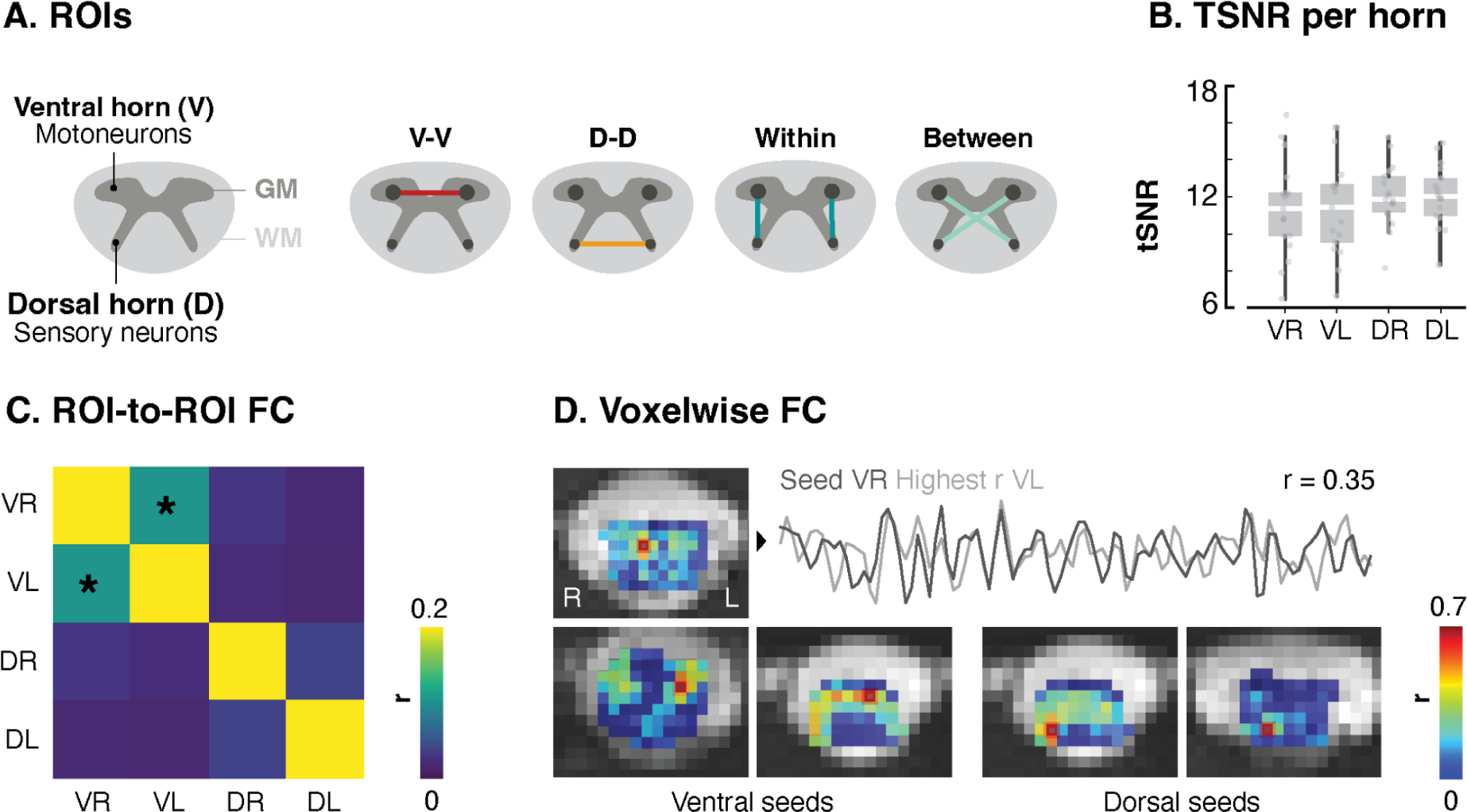
Extension of resting-state FC to the lumbosacral cord. A. Schematic cross-sections of the spinal cord. The left panel illustrates the spinal cord’s structure, featuring the characteristic butterfly-shaped gray matter (GM) surrounded by white matter (WM). The GM can be divided into four horns, housing motor (ventral horns, V) and sensory (dorsal horns, D) neurons. The right panel outlines potential connectivity patterns between these four regions of interest (ROIs). **B.** TSNR in the four ROIs, for the *PNM+Moco+CSF* denoising strategy. Each box represents the distribution (*i.e.*, from the 25th to the 75th percentile) of tSNR values across participants, with medians represented by the horizontal white line inside the box. Vertical lines correspond to the 1.5 interquartile range and dots represent tSNR values for each of the 17 participants. **C.** 4×4 correlation matrix showing FC between the four horns (seed-to-seed analysis, for the *PNM+Moco+CSF* denoising strategy). A significant connectivity is observed only between bilateral ventral horns (r = 0.11). **D.** Example slices showcasing results from the seed-to-voxels analysis, for the *PNM+Moco+CSF* denoising strategy. The seed voxel is highlighted, with resulting correlations (*i.e.*, other voxels) overlaid on the mean functional image. In the top row, correlation patterns for a seed in the left ventral (VR) horn are shown. The highest correlation is observed in the contralateral ventral horn (r = 0.35). Time courses of both seeds are presented. The bottom row presents additional connectivity maps for ventral seeds on the left panel, while examples for dorsal seeds are presented on the right panels. VR = ventral right, VL = ventral left, DR = dorsal right, DL = dorsal left.

Static functional connectivity was estimated by means of Pearson correlation coefficients and two types of analyses were conducted: i) seed-to-seed, and ii) seed-to-voxels.

#### 2.4.1. Seed-to-seed functional connectivity

Functional connectivity was computed using a region of interest (ROI)-based approach where we considered the 4 seeds to extract ROI-specific time courses (*i.e.*, average time courses for each seed and participant). This methodology aligns with earlier work in the cervical spinal cord (Barry et al., 2014; Eippert et al., 2017a; Kaptan et al., 2023; Kinany et al., 2019; Kong et al., 2014; Weber et al., 2016). We computed Pearson correlation coefficients between those time courses, resulting in a 4×4 matrix that summarizes the connectivity patterns of interest: ventral-ventral (VV), dorsal-dorsal (DD), within (W) and between (B) hemicords (Fig. 1A). We performed this analysis across all the denoising procedures to compare the impact of each strategy on functional connectivity estimates.

The significance of functional connectivity estimates was assessed using non-parametric tests. Indeed, even after applying a Fisher z-transformation to the correlation values, the Kolmogorov-Smirnov test rejected the assumption of normal distributions. Consequently, the Wilcoxon test was conducted on the correlation values to evaluate which connectivity pattern was significantly different from zero using the different denoising techniques (corrected for multiple comparison with Benjamini-Hochberg method).

#### 2.4.2. Seed-to-voxels functional connectivity

Functional connectivity was estimated within each slice, focusing on a single seed (*i.e.*, ROI) at a time. The correlations between the time courses of this ROI and those of each voxel within the slice were computed. This enabled visual assessment of the prevalence of specific patterns, similar to earlier work at 7T in the cervical spinal cord (Barry et al., 2014). For the sake of brevity, we present these results exclusively for time series processed using the denoising approach typically used in spinal cord fMRI (*i.e*., *PNM+Moco+CSF*).

#### 2.4.3. Temporal reliability and consistency

To investigate the temporal reliability of lumbosacral resting-state functional connectivity patterns, we split the fMRI time series of each participant into two halves in which correlation values were independently extracted. We computed the intraclass correlation coefficient (ICC) to measure the consistency of functional connectivity estimates across the temporal splits, for each denoising procedure. For this analysis, we used a two-way random effect model, ‘Case 2’ intraclass correlation coefficient defined as:

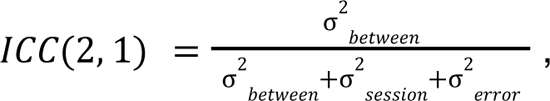

where σ^2^_between_ corresponds to the variance between participants and σ^2^_sessions_ indicates the variance between sessions (*i.e.*, the two halves). This metric, known as ‘absolute agreement’ in the literature, quantifies the proportion of total variance attributed to between-participants differences (Kaptan et al., 2023; McGraw and Wong, 1996; Shrout and Fleiss, 1979). To assess the uncertainty of this metric, we calculated the 95% confidence interval (CI) of the ICC values using a bootstrap procedure implemented in Python with the *pingouin* library (Vallat, 2018). According to established standards (Cicchetti and Sparrow, 1981; Hallgren, 2012), the ICC values are interpreted as follows: poor <0.4, fair 0.4-0.59, good 0.6-0.74, excellent ≥ 0.75.

In addition to investigating the test-retest reliability of functional connectivity estimates, we also calculated ICC in the split-half datasets for the following metrics: i) CSF, ii) cardiac and iii) respiratory. For i) CSF, we computed the mean amplitude of the signal from the power spectral density of each half of the regressor. For (ii) cardiac, we computed the average difference between cardiac peaks. For iii) respiratory, respiration traces were first band-pass filtered (cut-off frequencies: 0.01 Hz and 0.6 Hz), and median filtered over 1s. Subsequently, we identified the respiratory cycle by applying a Hilbert transform and by computing the phase of the signals (which measures the position of a waveform in time). We then determined the occurrence of zero-crossings within the detected respiratory cycles.

## 3. Results

### 3.1. Functional data quality control

Out of the initial 22 participants, 5 were excluded from the study due to excessive motion (*i.e.*, average FD > 0.4mm). In Figure 1B, the tSNR values for the remaining 17 participants are presented independently for each of the 4 horns (using the conventional *PNM+Moco+CSF* denoising scheme): ventral right = 11.33 (9.85-12.18) (median across participants and interquartile range, IQR), ventral left = 11.39 (9.51-12.64), dorsal right = 11.82 (11.12-13.07), dorsal left = 11.98 (10.94-12.98). Figure S1 includes an example tSNR map along with the corresponding mean functional image.

### 3.2. Extending resting-state FC to the lumbosacral spinal cord

A main goal of this study was to extend resting-state FC fMRI findings beyond the cervical spinal cord, by deploying such analyses at the lumbosacral level. To this end, we investigated connectivity patterns between the ventral and dorsal horns of both left and right hemicords (Figure 1C). Using this ROI-based approach, we observed a significant positive correlation between bilateral ventral horns (mean correlation value r = 0.11, Wilcoxon test, W = 24, p = 0.01, Benjamini-Hochberg corrected). When assessing the robustness of these connectivity estimates in each participant, we observed that ventral connectivity was positive in 70.6% of participants. Although no significant average correlation was reported between the other ROIs, 64.7% of participants exhibited positive bilateral dorsal connectivity, 64.7% positive dorsal-ventral within-hemicord connectivity, and 58.8% positive dorsal-ventral between-hemicord connectivity.

A seed-to-voxels analysis further highlighted these connectivity patterns. Figure 1D (top panel) illustrates a compelling example wherein a seed placed in the ventral region resulted in a strong correlation in the contralateral ventral horn. On average, among the participants, when the seed was placed in one of the two ventral horns, 51% of the slices exhibited a high positive correlation in the contralateral ventral horn. Notably, within this percentage, 26% displayed the distinctive butterfly shape characteristic of the gray matter (*i.e.*, FC between all four horns) (Figure 1D, bottom panel, left). Alternatively, when the seed was positioned in the dorsal horn, 47% of the slices showed a high correlation in the contralateral dorsal horn, with 20% of these displaying the gray matter butterfly shape (Figure 1D, bottom panel, right).

### 3.3. Impact of denoising strategies on signal properties

Given the inherent sensitivity of spinal cord fMRI to various noise sources (*e.g.*, breathing, heart rate, motion, …), we systematically and quantitatively compared the impact of different denoising techniques.

First, we assessed changes in signal quality, by evaluating tSNR and variance explained (*i.e.*, adjusted R^2^) for each applied strategy (Figure 2). We observed that the addition of nuisance regressors led to an increase of tSNR, with all denoising approaches significantly increasing the tSNR compared to the *baseline* pipeline (p < 0.001, Welch’s t-test corrected for multiple comparisons). The largest changes were observed when going from the *baseline* pipeline to mild denoising (*e.g.*, *CSF* or *Moco* pipelines, 24% and 24.5% increase compared to the baseline, respectively), and when adding the PNM-related regressors (*e.g.*, 31% for the *PNM* pipeline). The tSNR reaches its highest value for the most stringent denoising technique (*i.e.*, with all regressors combined) (11.85 (10.93-13.13), median across participants (IQR), 34% increase compared to *baseline*). A participant’s tSNR map example is provided in the supplementary material, depicted in Figure S1.

**Figure 2.**
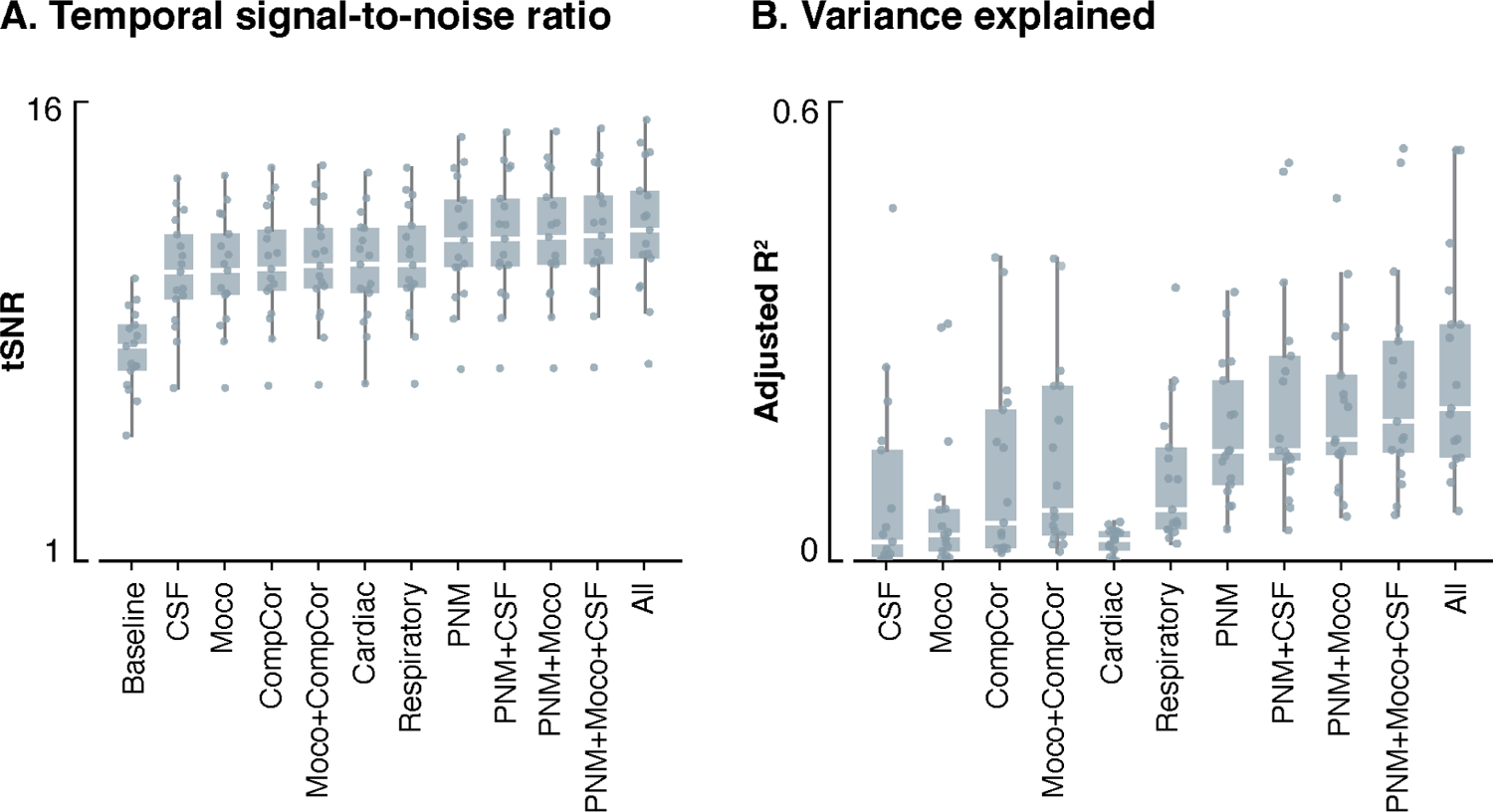
Signal properties following distinct denoising strategies. Temporal signal-to-noise ratio (tSNR) (**A.**) and variance explained (adjusted R^2^) (**B.**) values (y-axis) are presented for each denoising strategy (x-axis). The boxes represent the interquartile range (IQR), spanning from the 25th to the 75th percentile of the data, with the horizontal white line within each box indicating the median value across participants. Each dot represents the average metric (tSNR and adjusted R^2^) of the voxels within the spinal cord mask for a specific participant.

As for R^2^, the peak was also observed when combining all regressors. Generally, methods incorporating PNM regressors (*PNM*: 0.14 (0.10-0.23), *PNM+CSF:* 0.14 (0.13-0.27), s*PNM+Moco:* 0.16 (0.14-0.24), *PNM+CSF+Moco:* 0.18 (0.14-0.29), *PNM+CSF+Moco+CompCor:* 0.20 (0.13-0.31), median R^2^ across participants (IQR)) explained more variance than those who did not (*CSF:* 0.02 (0-0.14), *Moco:* 0.03 (0.01-0.07), *CompCor:* 0.05 (0.01-0.20), *Moco+CompCor:* 0.03 (0.03-0.23), *Cardiac:* 0.03 (0.01-0.04), *Respiratory:* 0.06 (0.04-0.15)). When comparing procedures sequentially, we observed that *Cardiac* exhibited a R^2^ significantly lower than the surrounding approaches (*Moco+PNM,* p < 0.05, *Respirator*y, p < 0.01, Welch’s t-test corrected for multiple comparisons).

### 3.4. Impact of denoising strategies on functional connectivity

We then focused on assessing the different denoising procedures from the perspective of functional connectivity (Figure 3). We consistently observed that only the connectivity between bilateral ventral horns (VV) appeared to be significant, regardless of the denoising method employed. Specifically, the highest connectivity values were obtained for the non-denoised time series (0.18 (0-0.25), median across participants (IQR)), as well as for the images denoised with the *Cardiac* pipeline (0.15 (0.01-0.34)). Stricter denoising procedures led to reduced functional connectivity, with VV connectivity being the weakest for the *All* pipeline (0.08 (0-0.18)).

**Figure 3.**
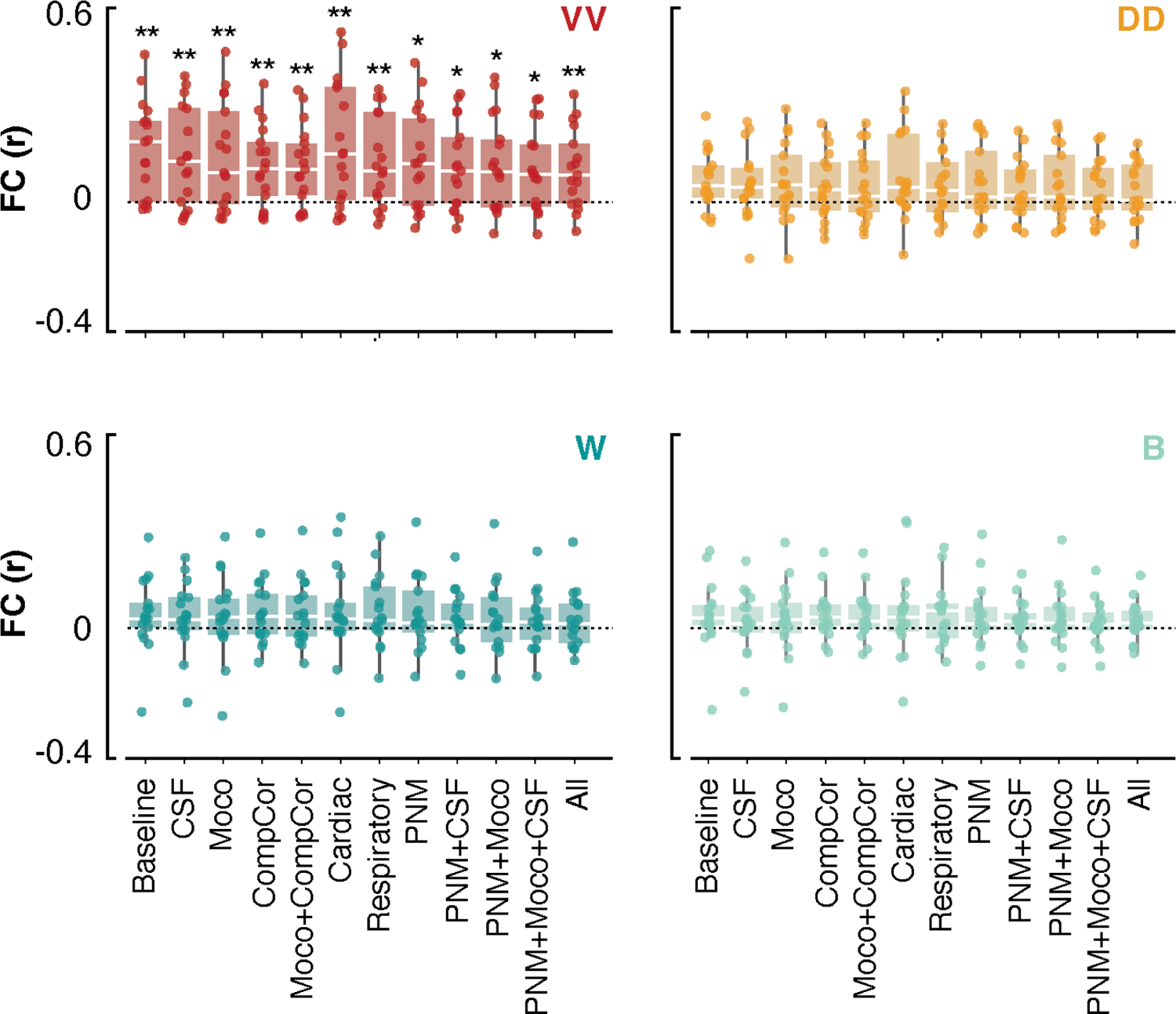
Static functional connectivity using distinct denoising strategies. For each seed-to-seed pattern (Figure 1A), we present functional connectivity estimates (y-axis) for each denoising strategy (x-axis). The boxes represent the interquartile range (IQR), spanning from the 25th to the 75th percentile, with the horizontal white line within each box indicating the median value across participants. Each dot represents the mean FC (across slices) for a specific participant. * indicates p < 0.05 and ** p < 0.01.

Conversely, the analysis of other connectivity patterns (DD, between and within horns connectivity) did not yield statistically significant results for any of the denoising procedures. However, DD exhibited higher median correlation values on average (0.05 ± 0.1) compared to between and within connectivity patterns (0.03± 0.08 and 0.04 ± 0.1, respectively). Further statistical testing confirmed the significance of DD connectivity being significantly higher than between (Wilcoxon test, p = 0.003, W=9441) and within (p = 0.017, W=10005) correlations.

### 3.5. Temporal reliability

Finally, we investigate the temporal reliability of the signals (Figure 4, see Figure S2 for scatter plots). In general, VV connectivity demonstrated a high reliability, predominantly within the good range (average ICC over denoising strategies = 0.67). Conversely, DD connectivity mostly fell within the poor range (average ICC = 0.49). For all connectivity patterns, a similar ICC profile was observed, albeit with different amplitudes. Notably, the *Cardiac* denoising strategy consistently exhibited the highest temporal reliability, achieving an excellent rating for VV (ICC = 0.83), within (0.81), and between (0.81) connections, and a good rating for DD (0.67). For VV, W, and B connectivity patterns, the strongest decline was observed when removing *CompCor* regressors, which brought reliability in the fair (VV and B) and poor (W) ranges. For DD, the removal of the *CSF* signal had the most impact, resulting in an ICC value in the poor range. On the stringent side of the denoising spectrum, we observed that the *PNM* (ICC = 0.79 for VV, 0.57 for DD, 0.76 for W, and 0.78 for B) and *PNM+Moco* (ICC = 0.73 for VV, 0.52 for DD, 0.72 for W and B) strategies exhibited the largest reliability.

**Figure 4.**
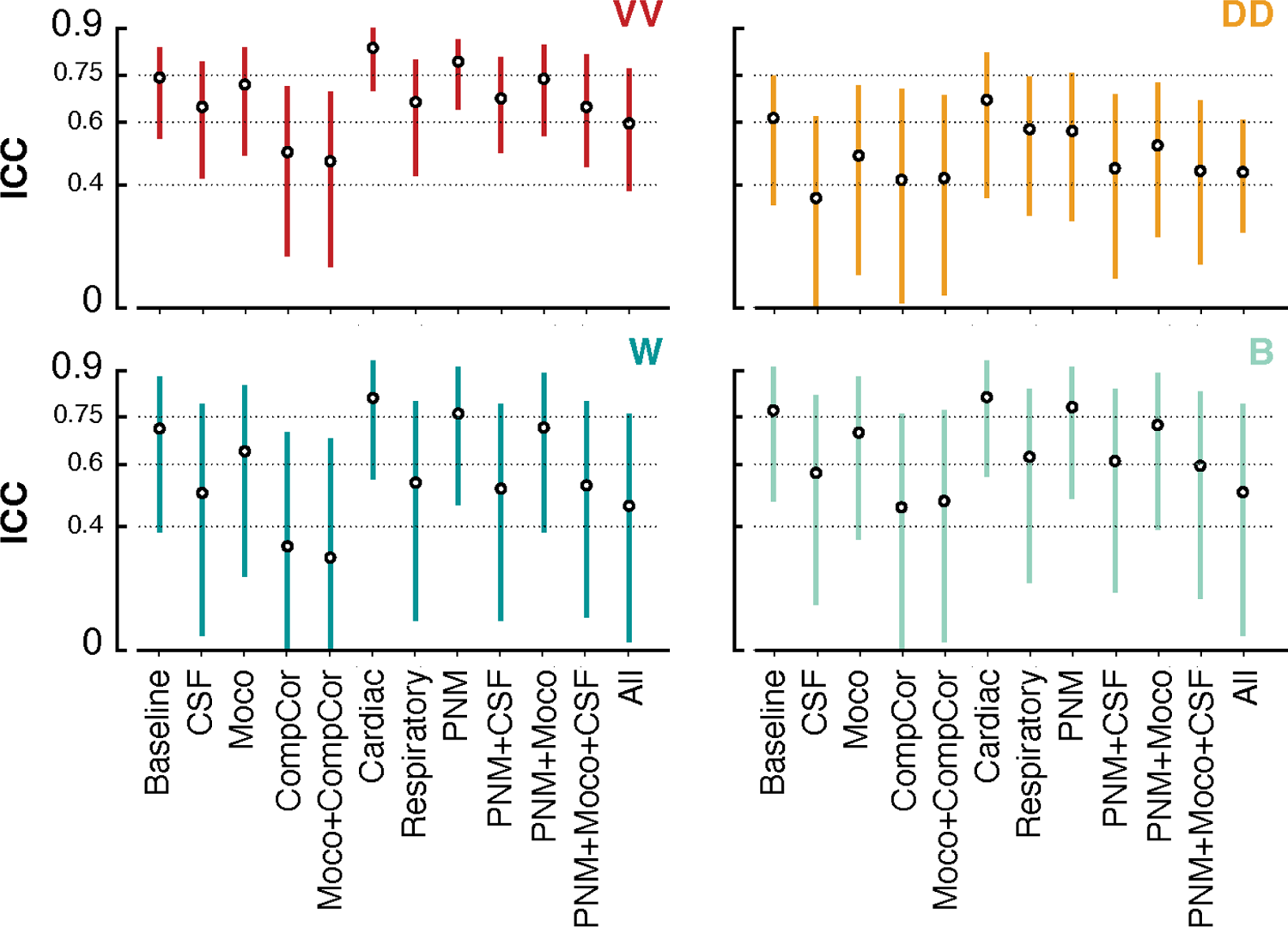
Temporal reliability using ICC. Distribution of ICC (Interclass Correlation) scores, focusing on the robustness and reliability of functional connectivity patterns derived from the fMRI signals of each participant. Each point in the figure represents the ICC score for a specific denoising technique employed in the analysis (x-axis) and the bars represent the confidence intervals at 95%. The dotted lines indicate the ICC ranges: poor < 0.4, fair 0.4-0.59, good 0.6-0.74, excellent ≥ 0.74.

To further elucidate the impact of removing specific noise signals on reliability, we computed ICC values for CSF, respiratory and cardiac time series, which all fell in the excellent range. Specifically, the CSF exhibited the highest ICC score with 0.95 (confidence interval at 95%, CI95% [0.87, 0.98]), followed by the cardiac signals with 0.94 (CI95% [0.87,0.97]), and finally the respiratory ICC value was 0.89 (CI95% [0.73, 0.96]).

## 4. Discussion

In recent years, a growing body of evidence has highlighted distinct spatial patterns of spontaneous activity within the human spinal cord at rest, consistently revealing correlations between its horns (Barry et al., 2018, 2014; Eippert et al., 2017b; Kaptan et al., 2023; Kinany, 2022; Kong et al., 2014). However, while these analyses have provided insights into the functional architecture of the cervical region of the spinal cord, a significant gap remains in the examination of such patterns within the lumbosacral area. In this study, we tackled this by systematically investigating horn-to-horn functional connectivity in the lumbosacral spinal cord of healthy participants. We first demonstrated that, akin to the cervical spinal cord, characteristic connectivity patterns can be identified in the lumbosacral region. In a subsequent step, we assessed the impact of different denoising strategies on these functional connectivity estimates.

### 4.1 Imaging the lumbosacral spinal cord

In spite of its relevance for healthy and impaired human behavior, the human lumbosacral spinal cord has been largely unexplored in neuroimaging studies. Previous studies examining lumbosacral activity were primarily conducted during task (Jia et al., 2019; Kornelsen and Stroman, 2007, 2004; Moffitt et al., 2005; Stroman et al., 2004), using a non-BOLD contrast mechanism known as signal enhancement from extravascular water protons (SEEP) (Stroman et al., 2001a, 2001b). However, the reliability of SEEP has been a matter of debate (Bouwman et al., 2008; Jochimsen et al., 2005). Only one recent study capitalized on the BOLD signal within the lumbosacral cord during resting-state scans, providing evidence of discernible patterns of functional connectivity in the spinal cord (Combes et al., 2023).

Several distinctions between cervical and lumbosacral imaging are noteworthy. One primary concern is the marked size difference. The cross-sectional dimensions and lengths of lumbosacral segments are notably smaller (7.7 ± 2.2 mm in average) compared to the more extensively studied cervical region (13.3 ± 2.2 mm) (Frostell et al., 2016; Kinany et al., 2022). Furthermore, the anatomical positioning of both regions implies the presence of different organs in their vicinity, potentially rendering them differentially susceptible to physiological noise. Interestingly, it has been suggested that lumbosacral regions are less prone to cardiac-related motion artifacts (Figley et al., 2008).

In addition to its smaller size, the lumbosacral cord is also characterized by high intersubject variability (Tins and Balain, 2016; Van Schoor et al., 2015), marked by substantial shifts between spinal segments and vertebrae, thus making conventional normalization based on vertebral landmarks suboptimal. To overcome these challenges, we opted to conduct all analyses in the native space of the participants, wherein we manually placed seeds in the different horns to serve as our regions of interest (ROIs).

### 4.2 Extension of static functional connectivity to the lumbosacral cord

In order to extend prior investigations of functional connectivity to the lumbosacral spinal cord, we resorted to established analysis techniques, commonly employed in the cervical spinal cord (Barry et al., 2018, 2014; Eippert et al., 2017b; Kaptan et al., 2023; Kinany, 2022; Kong et al., 2014). Through a ROI-based approach, we demonstrated significant functional connectivity between bilateral ventral horns. A subsequent slice-wise seed-to-voxels analysis, similar to the approach employed by (Barry et al., 2016), further underscored the presence of these bilateral motor networks. This corroborates the findings of Combes and colleagues (Combes et al., 2023), who similarly emphasized substantial connectivity between the ventral horns. However, it is noteworthy that the ventral connectivity reported in their study demonstrated an amplitude more than fourfold stronger than our observations. This discrepancy might be ascribed to the superior spatial resolution of their dataset, enhancing seed placement accuracy and mitigating partial volume effects. In addition, their use of trilinear interpolation, recognized for augmenting spatial smoothness in the dataset, could possibly contribute to the observed inflation in correlation values (Eippert et al., 2017a).

The observation of ventral-ventral connectivity echoes prior investigations in the cervical spinal cord, where it has been repeatedly documented, both in animal models (Chen et al., 2015; Wu et al., 2019, 2018), and in human studies that used various processing and acquisition procedures (Barry et al., 2018, 2014; Eippert et al., 2017b; Kaptan et al., 2023; Weber et al., 2018). These bilateral networks are postulated to arise from commissural interneurons connecting motoneurons from the two hemicords (Maxwell and Soteropoulos, 2020), serving multiple functions such as maintaining basal muscle tone – a state where motoneurons uphold posture and muscle tonicity even during quiescence (Latash and Zatsiorsky, 2015). In the lumbosacral spinal cord, which innervates lower limb muscles, these patterns may also be indicative of activity related to central pattern generators, pivotal in orchestrating locomotion (Jessell, 2009).

Notably, the ROI-based analysis did not reveal significant dorsal-dorsal connectivity, contrasting with results of Combes and colleagues (Combes et al., 2023). While dorsal networks have also been reported in several cervical studies (Barry et al., 2014; Eippert et al., 2017b; Kaptan et al., 2023), they were observed to be less pronounced (Barry et al., 2014) and less robust (Eippert et al., 2017b) than their ventral counterparts. Here, it should be noted that dorsal-dorsal patterns were still observed, both in a subset of participants (11 out of 17), as well as in the seed-to-voxels analyses. On one hand, this suggests that dorsal connectivity may be present but inherently more variable or weaker than the ventral connections. On the other hand, these disparities may partially be attributed to anatomical differences, given that dorsal horns are thinner than ventral ones. Consequently, seed placement for dorsal regions might be more sensitive to potential inaccuracies, which could influence the observed correlation estimates.

Finally, we did not observe significant connectivity either within (*i.e.*, dorso-ventral) or between (*i.e.*, left dorsal - right ventral and right dorsal - left ventral) hemicords. Such patterns have been observed in the lumbosacral (Combes et al., 2023) and cervical spinal cord (Eippert et al., 2017b; Kaptan et al., 2023; Weber et al., 2018). While these dorso-ventral connections may potentially support polysynaptic spinal reflexes (Sandrini et al., 2005), it is important to emphasize that they seem to be strongly influenced by the specific processing approach (Eippert et al., 2017a; Kaptan et al., 2023) and have exhibited poor reliability (Kaptan et al., 2023). Similar investigations in the lumbosacral spinal cord are essential to determine the reliability and physiological relevance of these connectivity patterns.

### 4.3 Impact of denoising on signal quality

Given that lumbosacral BOLD imaging has been virtually unexplored, a primary contribution of this study was the systematic evaluation of diverse denoising strategies on the time series. This comprehensive assessment included the examination of both variance explained (*i.e.*, adjusted R^2^) and their impact on the temporal signal-to-noise ratio (tSNR).

Notably, a progressive enhancement in tSNR was observed with more rigorous denoising. This trend was mirrored in the adjusted R^2^, with the exception of the *Cardiac* pipeline, which exhibited limited explanatory power, hinting at a restricted influence of cardiac physiological noise in this spinal cord region. We could argue that the lumbar region is relatively spared from the influence of cardiac signals due to its anatomical distance from the heart, shielding it from the pulsating movement. From an anatomical point of view, the heart overlaps more with the upper regions of the spinal cord, and its upward beating within the respiratory cage may have a more pronounced impact on the cervical region compared to the lumbar region. These considerations are in line with work reporting limited cardiac-related motion in the caudal part of the spinal cord (Figley et al., 2008).

Of particular interest was the pivotal role of *PNM* regressors in improving the tSNR, underscoring their efficacy in capturing the variance of the signal. In contrast, using *CompCor* regressors to account for physiological noise yielded a more moderate effect on signal quality. These results suggest that accounting for the interaction between cardiac and respiratory signals is valuable. Besides, it hints at the fact that, despite the position of the lumbosacral region, relatively distant from the heart and lungs, mitigating potential physiological signals remains beneficial. Indeed, even in the brain, physiological fluctuations can induce notable change in fMRI time series, shown to lead to “physiological networks” (Chen et al., 2020) reminiscent of large-scale networks conventionally attributed to distantly synchronized neuronal activity.

In light of these results, our recommendation is to adopt a denoising pipeline that incorporates PNM regressors to achieve optimal enhancement of signal quality. This is in agreement with observations in the cervical spinal cord (Brooks et al., 2008; Kong et al., 2012), where PNM was found to be an effective denoising strategy, notably by eliminating false-positive activations, such as active voxels in the CSF space surrounding the cord.

### 4.4 Robustness and reliability of functional connectivity

Since our work primarily centered on functional connectivity, our subsequent objective was to evaluate the extent to which the strength and reliability of connectivity patterns were influenced by the denoising procedures applied.

While our analyses revealed a general decrease in functional connectivity with more stringent denoising, bilateral ventral networks appeared to be significant regardless of the deployed denoising. This supports the genuine nature of these motor networks, emphasizing their robustness. Conversely, none of the other connectivity patterns reached significance in any of the conditions.

To further explore the robustness of these connectivity estimates, we deployed a split-half analysis to evaluate their reliability and reproducibility by means of intraclass correlation coefficients (ICC). Reassuringly, VV connections appeared to be the most reliable, with ICC values primarily scoring as good (ICCs > 0.6). In comparison, cervical networks obtained using correlation analyses have been shown to exhibit a fair to good level of reproducibility, both at 7T (Barry et al., 2016) and 3T (Kaptan et al., 2023) field strengths. Likewise, similar levels of reliability were reported for networks retrieved using independent component analysis (Kong et al., 2014).

Upon closer examination of the relationship between denoising strategies and the reliability of functional connectivity, we did not observe a consistent decrease in reliability with an increasing number of regressors, unlike recent findings in the cervical spinal cord (Kaptan et al., 2023). Instead, we noted that the removal of CSF-related regressors, as seen in both the simple *CSF* pipeline or the *CompCor* approach, had the most significant impact on reliability. We posit that the sensitivity of the lumbosacral signals to CSF removal may pertain to the substantial volume of the subarachnoid space in these segments. Specifically, the relative average area of the spinal cord in relation to CSF in the lumbosacral cord is 29.4%, as opposed to 52.5% in the cervical region (percentages determined by calculating the average cross-sectional area of the PAM50 masks at the respective levels). This may imply additional CSF-related motion in the caudal region of the cord. Provided that CSF signals seem to exhibit structured properties, as evidenced by their ICC score in the excellent range, the CSF pipeline may impact the reliability by removing a large portion of reliable artifactual signals. In comparison, even though they also demonstrated high reliability, physiological signals – particularly the cardiac ones – may have a comparatively lesser impact on FC reliability, owing to their more limited influence on these segments of the cord.

Meanwhile, we observed that PNM-based pipelines led to robust patterns of connectivity, achieving ratings from good to excellent for VV connections. Considering the substantial portion of variance removed by these denoising techniques (as indicated by the adjusted R^2^), this suggests that they may be a good approach to eliminate nuisance signals while preserving meaningful functional connectivity. Notably, even when combined with CSF regressors (*PNM+CSF* or the standard *PNM+CSF+Moco*), VV connectivity remained within the good range.

### 4.5 Limitations of the study

The current study has several limitations that warrant acknowledgment. First, a major constraint is inherent in the manual placement of the seeds, constituting a potential source of variability. This method may have resulted in underestimated connectivity estimates, particularly for the dorsal horns, where seed placement can be challenging due to their elongated geometry. Second, the reliability estimates were derived from split-half time series, potentially leading to inflated values compared to those obtained from separate test-retest runs, despite the length of the runs (15 min in total). Employing distinct test-retest sessions could offer a more accurate reflection of the reliability of functional connectivity patterns.

### 4.6 Conclusion and outlook

In summary, our findings underscore the existence of intrinsic functional connectivity in the lumbosacral region, in the form of bilateral ventral connectivity. Importantly, the consistency and robustness of these connectivity patterns were confirmed by their persistence across various denoising strategies. In addition, our results hint at the effectiveness of physiological noise modeling (PNM) as a valuable approach for denoising lumbosacral spinal cord fMRI images, while preserving the strength and reliability of functional connectivity estimates. Finally, given the nascent stage of lumbosacral fMRI research, future investigations are needed to probe these findings across diverse acquisition and processing schemes. While the current study proposes a first step in this direction, further research is necessary to ascertain the robustness and broader applicability of the observed functional connectivity patterns in the lumbosacral spinal cord.

## Data and Code Availability

Code is publicly available on GitHub (https://github.com/iricchi/Lumbar.git). The data can be accessed on Mendeley Data with the identifier doi: [to be done upon acceptance]

## Authors contributions

I.R. N.K., and D.V.d.V. initiated the study and wrote the paper. N.K. designed the protocol and I.R. and N.K. collected the data. I.R. processed and analyzed the data.

## Supplementary material

**Figure S1.**
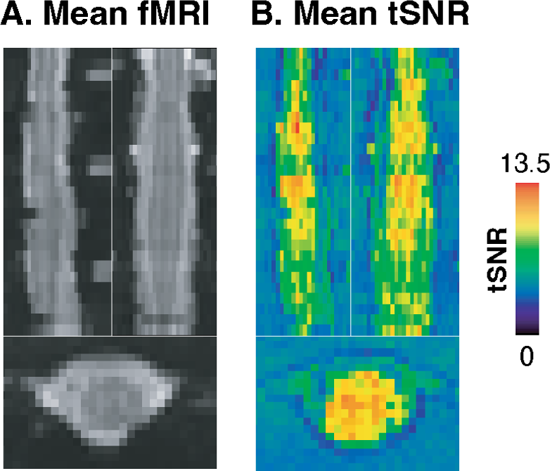
Data quality specific subject. (A) Mean functional image for an example participant. Sagittal, coronal, and axial views are shown. (B) Corresponding tSNR map

**Figure S2.**
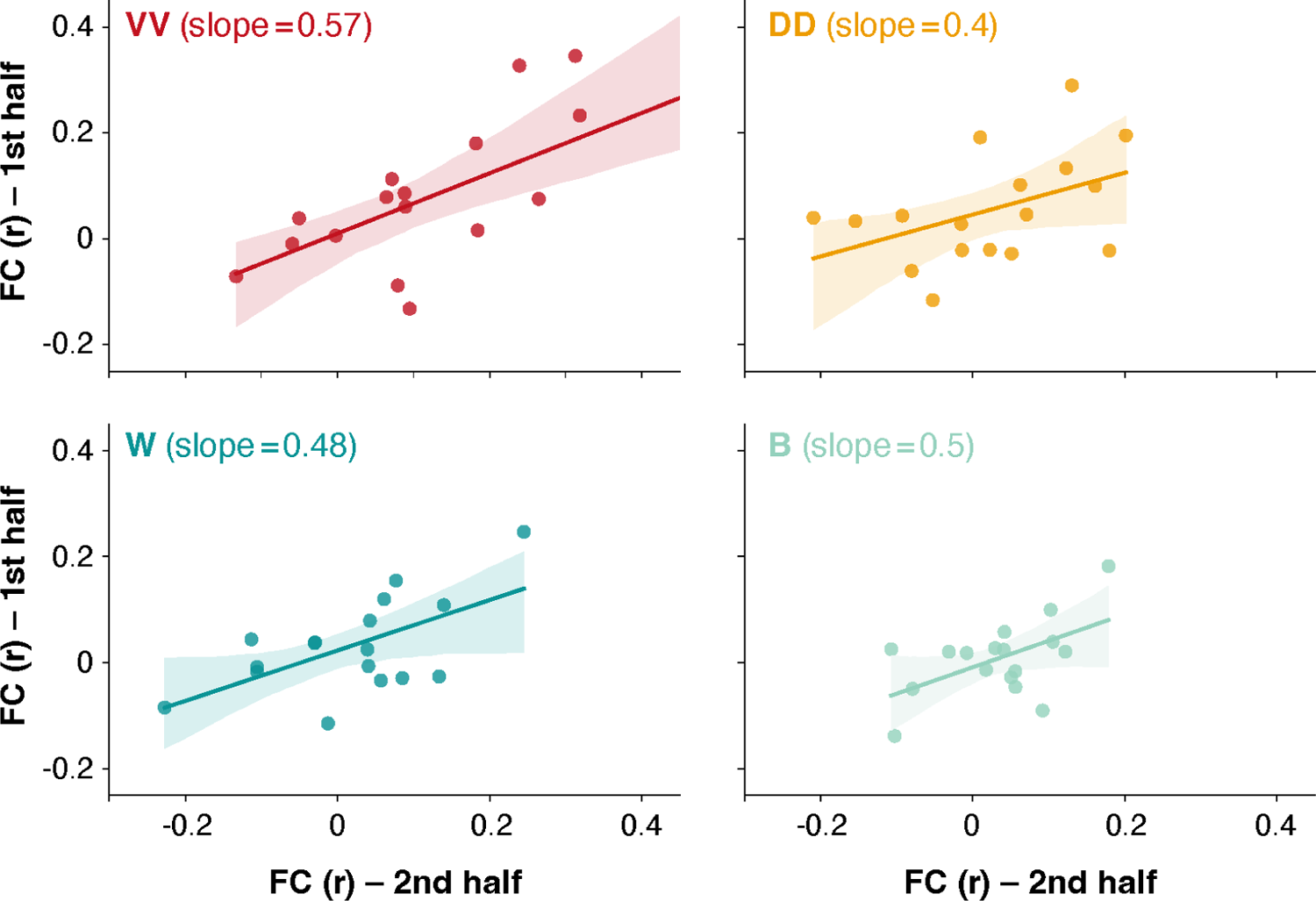
FC temporal stability. Scatter plots illustrate the correlation values in each half dataset plotted against each other, with the x-axis representing the first half and the y-axis representing the second half). Correlations values are obtained using the time courses denoised with the *PNM+Moco+CSF* pipeline. Ventral-ventral (VV) connectivity is depicted in red, dorsal-dorsal (DD) in yellow, within horns in dark green, and between horns in light green. The slopes for VV, DD, within horns (W), and between horns (B) are 0.57, 0.4, 0.48, and 0.5, respectively.

